# Azole resistance: Patterns of amino acid substitutions in *Candida* sterol 14α-demethylase

**DOI:** 10.1101/2024.07.04.602050

**Authors:** R. Shyama Prasad Rao, Larina Pinto, Rajesh P. Shastry, Tikam Chand Dakal, Prashanth N. Suravajhala, V. K. Sashindran, Sudeep D. Ghate

**Author notes:** School of Biological Sciences, University of Edinburgh, United Kingdom (Larina Pinto).

## Abstract

The emergence of azole-resistant *Candida* infections is a major concern. A key mechanism is the gain of resistance through amino acid substitutions in the sterol 14α-demethylase, the main target of azole drugs. While numerous resistant substitutions are known, the pattern of such substitutions is unclear. We hypothesized that the resistant substitutions occur disproportionately at the azole-binding sites. We compiled 2,222 instances of azole-resistant substitutions from the literature and performed extensive computational sequence analyses. Altogether there were 169 known substitutions at 133 sites in sterol 14α-demethylases of seven *Candida* species, whereas *C. albicans* alone had 120 substitutions at 97 sites. Just 10 sites and 18 substitutions (such as Y132F/H, K143R, D116E, and G464S) accounted for 75% of the total instances. Only about 48% of the sites were present within the previously recognized hotspot regions, while just 33% of the known azole-interacting residues had known resistant substitutions, most of them with only a few instances. The literature data on azole-resistant substitutions in *Candida* appear to be highly biased as a few substitutions such as Y132F/H and K143R were preferentially sought and reported with over 1000 instances, while there were also numerous reports of “resistant” substitutions in azole-susceptible *Candida* isolates. Our study provides interesting perspectives into azole resistance.

## 1. Introduction

Fungi cause considerable human morbidity and mortality, and are responsible for at least 13 million infections and 1.5 million deaths per year globally (Bongomin et al., 2017; Pfaller and Diekema, 2007; Vallabhaneni et al., 2016). The burden is underappreciated in different countries; but in populous countries such as in India an estimated 4.1% of the population suffer from serious fungal infections (Ray et al., 2022) and in China at least 5% of the population is at risk (Zhou et al., 2020). Invasive candidiasis, infections caused by the yeast *Candida*, is on the rise globally, with an estimated incidence of 3-5 per 100,000 persons in the general population (Pappas et al., 2018).

Women have an increased burden. About 75% of women develop vulvovaginal candidiasis at least once in their lifetime. Worldwide, recurrent vulvovaginal candidiasis affects about 138 million women annually (Denning et al., 2018).

The most common *Candida* sp. associated with invasive candidiasis is *C. albicans* (Pappas et al., 2018; Pfaller and Diekema, 2007). However, many other species, such as *C. auris, C. glabrata, C. parapsilosis*, and *C. tropicalis*, are increasingly being isolated from clinical samples. Some reasons for this include immune-compromised states secondary to chemotherapy, bone marrow/stem cell transplantation, and solid organ transplantation. The fungal isolate profiles in solid-organ transplant patients are similar to that in non-transplant patients with *C. albicans* accounting for 50% of the invasive candidiasis. The other species detected include *C. glabrata, C. parapsilosis*, and *C. tropicalis*. But stem-cell transplant patients are at greater risk for non-albicans *Candida* spp. with *C. krusei* and *C. guillermondii* being more common (Marr et al., 2000; Pappas et al., 2010; Pfaller and Diekema, 2007). Four major classes of drugs, namely azoles, echinocandins, polyenes, and flucytosine are used for treating fungal infections (Arendrup and Patterson, 2017). Azole antifungals such as fluconazole are frequently used to treat *Candida* infections as they are inexpensive, have limited toxicity, and can be orally administered. While most *C. albicans, C. parapsilosis*, and *C. tropicalis* are sensitive to fluconazole and other drugs, there are reports of isolates resistant to fluconazole. *C. krusei*, in particular, shows resistance to fluconazole and variable sensitivity to itraconazole, 5FC, and amphotericin B. (Pappas and Silveira, 2009; Pristov and Ghannoum, 2019; Whaley et al., 2017). The drug-resistant *Candida* infections are also common in immunocompromised individuals such as in cancer patients (Farmakiotis et al., 2014). More alarmingly, multidrug resistant *Candida* which cause serious invasive infections with high mortality are increasingly being reported (Forsberg et al., 2019).

The key target of azole antifungals is the enzyme sterol 14α-demethylase (also called ERG11 or CYP51 protein in the cytochrome P450 family, encoded by *ERG11* gene) which catalyses the synthesis of ergosterol, a component of cell membrane, by removing the methyl group at C14 of lanosterol (Hargrove et al., 2017). By limiting the lanosterol binding to the enzyme, azoles inhibit the biosynthesis of ergosterol and thus interfere with the integrity of the fungal cell membrane. The minimum inhibitory concentration (MIC) breakpoint for fluconazole susceptible *C. albicans* isolates is ≤2 µg/ml and for resistant one is ≥8 µg/ml. In case of voriconazole, it is ≤0.12 µg/ml and ≥1 µg/ml, respectively (Pristov and Ghannoum, 2019). Consequently, one important mechanism of azole resistance is the emergence of numerous mutations (= amino acid substitutions in protein sequence) in sterol 14α-demethylase. In fact, such substitutions are widely reported in the clinical isolates of *Candida* (Xiang et al., 2013) and other fungi (Zhang et al., 2019). For example, Y132F/H is one of the most common substitutions, and isolates might have multiple substitutions simultaneously (Flowers et al., 2015; Rao et al., 2024; Xiang et al., 2013). Substitutions, individually or in combination of two or more, strongly alter the azole MIC (Flowers et al., 2015; Warrilow et al., 2019). Further, MIC might vary vastly between different substitutions and azoles.

For example, Y132F has a MIC of 64 to >128 µg/ml for fluconazole and 4 to >16 µg/ml for voriconazole, whereas K143R has a MIC of 64 to >128 µg/ml for fluconazole, but only 0.5 to 1 µg/ml to voriconazole (Healey et al., 2018). Similarly, different substitutions in same position might have different effects (Rao et al., 2024). For example, Y132F has a far higher MIC (>64 µg/ml) compared to Y132H (16 to 32 µg/ml) for fluconazole (Xiang et al., 2013).

Mechanistically, substitutions disrupt the interactions of ligands/azoles with the protein, and lower their binding efficacy (Rao et al., 2024). For example, many resistant (MIC: 16-256 µg/ml) clinical isolates of *C. tropicalis* were found to have two non-synonymous mutations A395T (Y132F) and C461T (S154F) in ERG11 gene (Paul et al., 2022). Using molecular docking, binding energies of fluconazole and voriconazole against the native protein were found to be -6.83 and -7.44 kcal/mol, respectively; whereas against the mutant protein they were -6.38 and -7.22 kcal/mol, respectively. Thus, there was 0.22 to 0.45 kcal/mol difference in the binding energies.

While numerous resistant substitutions in *Candida* sterol 14α-demethylase are known, the pattern of such substitutions is unclear as they have not been systematically explored thus far. There are several interesting open questions. For example, how frequent the different azole-resistant substitutions are? Do they have similar patterns of occurrence in different *Candida* species? And, do they preferentially occur in the substrate-binding sites (Rosam et al., 2021)? We hypothesized that the resistant substitutions occur disproportionately at the azole-binding sites. We sought to answer this by performing computational sequence analyses of 2,222 instances of azole-resistant substitutions collected from the literature. Our results provide valuable insights into azole resistance, and antifungal drug discovery and optimization.

## 2. Materials and Methods

### 2.1. Data curation on azole-resistant substitutions in sterol 14α-demethylase

We searched the databases – PubMed/Medline, Scopus, Web of Science, and Google Scholar – using various combinations of the following keywords: “*Candida*”, “azole resistance”, “mutation/substitution”, AND/OR “sterol 14α-demethylase”, and sifted through the research papers to find the reports of experimentally confirmed azole-resistant substitutions in various clinical isolates (Rao et al., 2024). The review reports citing substitutions, if any, were traced to the original reports. We found (as of Apr 24, 2024) at least 104 papers reporting a total of 2,665 instances of amino acid substitutions in the sterol 14α-demethylase from seven *Candida* species (Table S1).

### 2.2. Acquisition of sequence and structural information

The sterol 14α-demethylase protein sequences from seven *Candida* species (*C. albicans, C. auris, C. glabrata, C. krusei* – currently renamed as *Pichia kudriavzevii, C. orthopsilosis, C. parapsilosis*, and *C. tropicalis*) were downloaded from the UniProt website (https://www.uniprot.org/, last accessed on Apr 24, 2024) and the corresponding nucleotide sequences were downloaded from the NCBI website (https://www.ncbi.nlm.nih.gov/, last accessed on Apr 24, 2024).

The data on probable azole-interacting residues (that are well within a distance of 5 Å from the ligand; Mullins et al., 2011) were curated (Table S2) from multiple sterol 14α-demethylase 3D structures that are available in the Protein Data Bank (PDB, https://www.rcsb.org/, last accessed on Apr 24, 2024). However, the ligand-complexed protein 3D structures were present only for a few azoles and/or *Candida* species; and thus, information from other species such as from *Saccharomyces cerevisiae* were also used when available.

### 2.3. Sequence/data analyses

The number of sites with substitutions and their instances were enumerated for each sequence/species. The *Candida* sterol 14α-demethylase protein sequences were aligned (Fig. S1) using multiple sequence alignment web server Clustal Omega (https://www.ebi.ac.uk/Tools/msa/clustalo/, last accessed on Apr 24, 2024) for comparison of the substitution sites among the sequences/species. Each substitution was mapped to species-specific sterol 14α-demethylase protein sequence, and also categorized if it belonged to azole-binding sites and/or previously known hotspot regions (Marichal et al., 1999). The substitutions sites were mapped to the 3D structure of *C. albicans* sterol 14α-demethylase (PDB:5TZ1) and visualized using Visual Molecular Dynamics (VMD) molecular graphics software.

Functional impact of substitutions was assessed by using the evolutionary analysis of coding SNP (cSNP) tool in Protein ANalysis THrough Evolutionary Relationships (PANTHER) web server (https://pantherdb.org/, last accessed on May 30, 2024) (Thomas et al., 2003). It is based on substitution position-specific evolutionary conservation (subPSEC) score – a p < 0.05 means the substitution has no functional adverse effect (Silva et al., 2016; Thomas et al., 2003).

### 2.4. Statistical analyses

A Z-test for two proportions was performed using the prop.test function in R to test whether the proportions were significantly different (p < 0.05, two-tailed) from the expected.

## 3. Results

### 3.1. Azole-resistant substitutions in Candida sterol 14α-demethylase

After extensive search, we curated as many as 2,665 instances of amino acid substitutions that have been reported in 104 publications for sterol 14α-demethylase of seven *Candida* species (Table S1). The substitutions were mapped on to sterol 14α-demethylase protein sequences and compared among seven *Candida* species by using multiple sequence alignment (Fig. S1). However, many instances were problematic for various reasons such as synonymous mutations (Silva et al., 2016), issues in site mapping to the protein sequence, and/or their presence in the sensitive phenotype. After ignoring them, there were 2,222 instances of azole-resistant substitutions from 110 publications (Table 1). As many as 27 sites (20.3%) had multiple types of substitutions – for example, Y132 had three (Y132C, Y132F, and Y132H) and K143 had four (K143E, K143N, K143Q, and K143R) types (Table 2). Altogether there were 169 known substitutions at 133 sites in sterol 14α-demethylases of seven *Candida* species (Table 1). The *C. albicans* alone had 800 instances from 120 substitutions at 97 sites, whereas *C. parapsilosis* had the next highest of 729 instances from 12 substitutions at 12 sites (Table 1).

**Table 1.** Instances of azole-resistant substitutions in sterol 14α-demethylase of different *Candida* species.

| Species | Protein ID<sup>1</sup> | Seq. length | # of sites | # of substitutions<sup>2</sup> | # of instances<sup>3</sup> | # of studies |
| --- | --- | --- | --- | --- | --- | --- |
| *C. albicans* | P10613 | 528 | 97 | 120 | 800 | 55 |
| *C. auris* | A0A2H1A309 | 524 | 12 | 12 | 254 | 14 |
| *C. glabrata* | P50859 | 533 | 28 | 28 | 37 | 2 |
| *C. krusei*<sup>4</sup> | Q09GQ6 | 528 | 2 | 2 | 3 | 1 |
| *C. orthopsilosis* | H8X874 | 522 | 6 | 6 | 78 | 1 |
| *C. parapsilosis* | C7EXA5 | 522 | 12 | 12 | 729 | 30 |
| *C. tropicalis* | P14263 | 528 | 7 | 8 | 321 | 13 |
| Total<sup>5</sup> | - | 545 | 133 | 169 | 2,222 | 110<sup>6</sup> |

<sup>1</sup>From UniProt (https://www.uniprot.org/); <sup>2</sup>Each site/position may have more than one type of amino acid substitution;
<sup>3</sup>Based on all clinical isolates or sequences reported in various papers (see Table S1); <sup>4</sup>Currently renamed as *Pichia*
*kudriavzevii*; <sup>5</sup>Based on alignment of sequences from all seven species (see Fig. S1 for multiple sequence alignment);
<sup>6</sup>Unique studies.

**Table 2.** List of top-10 azole-resistant substitution sites (that together makeup ≈75% of the total instances) in *Candida* sterol 14α-demethylase.

| # | Site<sup>1</sup> | Amino acid | Substitutions | # of instances | # of studies |
| --- | --- | --- | --- | --- | --- |
| 1 | 132<sup>#</sup> | **Y** | **F, H, C** | 843 (774, 68, and 1) | 75 |
| 2 | 143 | **K** | **R, Q, E, N** | 176 (168, 5, 2, and 1) | 34 |
| 3 | 154 | **S**<sup>^</sup> | **F** | 149 | 9 |
| 4 | 398 | **r**<sup>^</sup> | **I, G** | 107 (106 and 1) | 12 |
| 5 | 116 | **D** | **E** | 91 | 18 |
| 6 | 464 | **G** | **S, D** | 84 (83 and 1) | 21 |
| 7 | 266 | **E** | **D** | 66 | 17 |
| 8 | 128 | **K** | **T, N** | 60 (59 and 1) | 15 |
| 9 | 257 | **Y** | **H** | 48 | 11 |
| 10 | 126<sup>#</sup> | **F** | **L** | 41 | 8 |

<sup>1</sup>Site index is based on *C. albicans* sequence; <sup>#</sup>Only Y132 and F126 are the azole-binding sites among the top-10 azole-resistant sites; <sup>^</sup>Only reported in non-*C. albicans*.

### 3.2. Pattern of azole-resistant substitutions

Except for Y132, K143, and a few other sites, majority (82.7%) of the substitutions/sites were identified only from one *Candida* species (Fig. 1A). A bar plot of the number of instances of substitutions at each site is shown in Fig. 1B. When present, they ranged from one to 843 instances per site. Interestingly, just 10 sites and 18 substitutions (such as Y132F/H, K143R, D116E, and G464S) accounted for 74.9% of the total instances (Table 2). With 774 instances, Y132F was the most common substitution reported. Although it was reported from six *Candida* species, most instances were from *C. parapsilosis* (506) and *C. tropicalis* (151). The substitution Y132H (67 instances) was more common than Y132F (21 instances) in *C. albicans*.

**Fig. 1.**
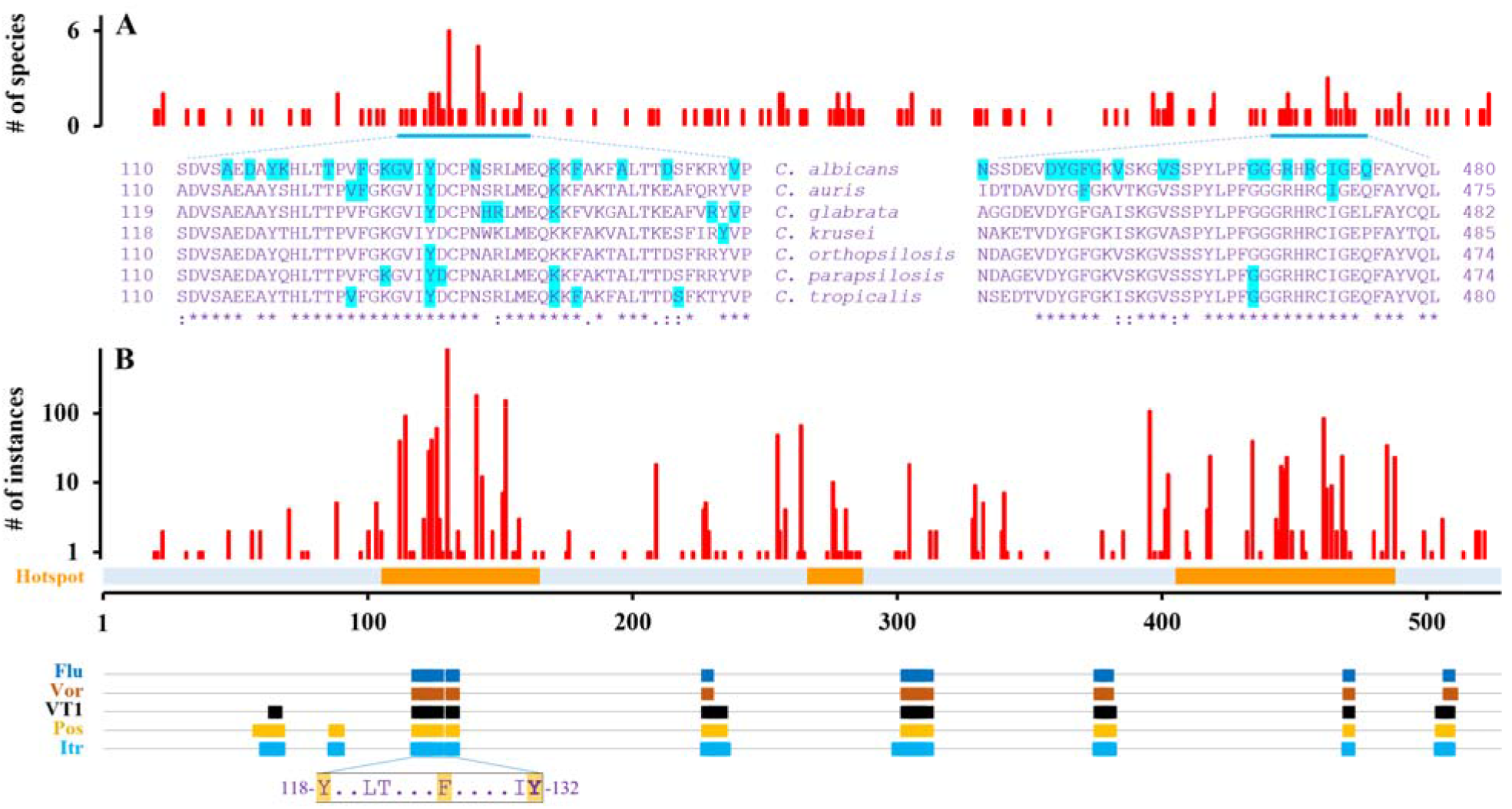
Substitution pattern in *Candida* sterol 14α-demethylase. (A) The bar plot shows the number of *Candida* species with known azole-resistant substitutions along the sterol 14α-demethylase. Multiple sequence alignments for two regions at 110-160 and 440-480 are shown with known substitution sites blue shaded. (B) The vertical bar plot shows the number of instances of substitutions at each site. There was a total of 2,222 instances at 133 sites, ranging from one to 843 instances per site. The three hotspot regions (based on Marichal et al., 1999) are shown as orange bars over the scale indicating the amino acid sequence positions (1 to 528, based on *C. albicans*). Lower horizontal bar plot shows the azole-interacting amino acid positions in sterol 14α-demethylase (based on PDB structures) for short-tail (fluconazole and voriconazole), medium-tail (VT1), and long-tail (posaconazole and itraconazole) azoles. Azole-interacting amino acids are shown for sequence region 118-132 in which three sites are known to have azole-resistant substitutions (highlighted in orange).

A previous study (Marichal et al., 1999) observed that the 98 instances of substitutions at 29 sites were not randomly distributed but clustered in three regions, and recognized them as three hotspot regions (amino acids 105 to 165, 266 to 287, and 405 to 488) in *C. albicans* sterol 14α-demethylase. In this study, we found that while 84.8% of the instances were present within, some 52.6% of the substitution sites were present outside of these previously recognized three hotspot regions (Fig. 1B).

Although many of the substitutions such as D116E, K143R, E266D, and V437I seemed to be conservative (Table S1); the functional analysis using cSNP tool indicated that only three substitutions – D116E and K128T/N – to be probably benign while others as probably damaging. Neither D116 nor K128 were azole-binding sites. While there were many instances of multiple substitutions (such as Y132F and K143R and Y132F and G307A) occurring in the same sequence (Arastehfar et al., 2020a), such information was not clear for most others sites/pairs.

### 3.3. Azole-binding sites versus resistant-substitutions

We gathered information on azole-interacting residues (that are within a distance of 5 Å from the ligand; Mullins et al., 2011) from multiple sterol 14α-demethylase 3D structures that are available in the Protein Data Bank (Table S2). As many as 18 residues interacted with short-tailed azoles (fluconazole and voriconazole) whereas 36 residues interacted with medium/long-tailed azoles (VT1, posaconazole, and itraconazole) (Table S2 and Fig. 1B). However, many sites such as F58, V234, L300, and V509 only infrequently interacted. Based on the overlap between the two sets, just 33.3% of the known azole-binding sites had known resistant substitutions (Fig. 2A). It was not significantly different (p = 0.435 for short-tailed azoles and p = 0.279 for long-tailed azoles; Z-test for two proportions, two-tailed) compared to the expected proportion of 25.2% (133 known substitution sites out of 528 positions). Further, only a few resistant substitutions/instances were known for most of those overlapping sites.

**Fig. 2.**
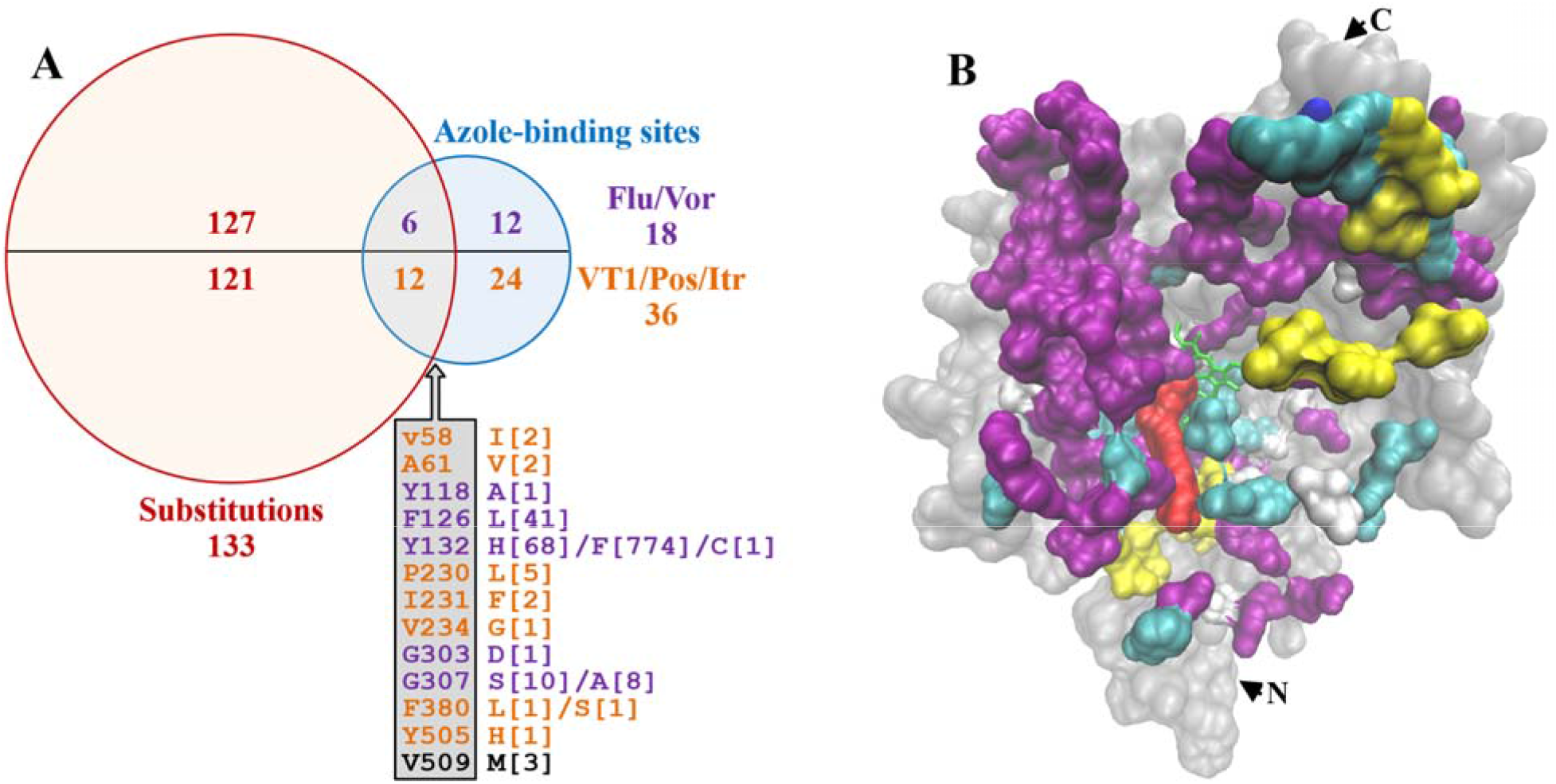
Azole-binding sites and resistant substitutions. (A) Venn diagram shows the overlap between the azole-binding sites and resistant substitutions. Only 33.3% of the azole-binding sites (6/18 for short-tailed and 12/36 for medium/long-tailed azoles) have known resistant substitutions (listed in the box below the intersection – sites shown in violet and black colour for sort-tailed, violet and orange for medium/long-tailed azoles, and the numbers in the square brackets are the instances). (B) The 133 known substitutions were mapped to *C. albicans* sterol 14α-demethylase 3D structure (PDB:5TZ1). Most of the substitutions are present far away from the ligand (red space fill), or on the surface of the protein (substitutions are shown as space fills – purple in α-helix regions, yellow in β-sheet regions, and cyan/white in loop regions. Overall protein shape is shown in grey space fill. The N- and C-termini are indicated by arrowheads).

The 133 known substitution sites were mapped to *C. albicans* sterol 14α-demethylase 3D structure (PDB:5TZ1). It was evident that most of the substitutions were present far away from the ligand binding pocket, and in fact, numerous sites were on the surface of the protein (Fig. 2B).

### 3.4. Substitutions in sensitive isolates

There were at least 26 instances of substitutions (at 17 sites) identified only in sensitive/susceptible isolates (Table S1). For example, substitutions such as D225H/Y, D247G, A255V, and N283Y were present in sensitive isolates (Arastehfar et al., 2020b; Asadzadeh et al., 2017; Ying et al., 2013), but not reported in resistant isolates. However, as many as 258 instances of known “resistant” substitutions (at 29 sites) were also reported in sensitive isolates (Table S1 and S3). For example, the “resistant” substitutions such as D116E, K128T, Y132F/H, E266D, and V437I were also present in sensitive isolates (Liu et al., 2015; Xiang et al., 2013; Ying et al., 2013).

## 4. Discussion

Antifungal resistance is a key global challenge and azole-resistant *Candida* infections are becoming a public health concern (Castanheira et al., 2020; Pfaller et al., 2019). Azole-resistance has been primarily linked, among other things, to substitutions in the sterol 14α-demethylase (also known as ERG11 protein). As a result, they have been keenly explored and numerous resistant substitutions were reported (Fan et al., 2023; Marichal et al., 1999; Morio et al., 2010; Xiang et al., 2013). However, the extent of such substitutions – for example, the number of sites and the frequency of substitutions at each site – was not obvious till date.

This work curated a large number of azole-resistant substitutions occurring in *Candida* sterol 14α-demethylase. It will become a useful catalogue for such studies in the future. Further, it provides insightful patterns – for example, the number and abundance of substitutions in each *Candida* species. It may be noted that when there were multiple substitutions, they came from different clinical/experimental isolates and/or from different studies, and therefore provided an independent validation of such resistant substitutions/sites (Rao et al., 2024).

A few sites such as Y132F and K143R were reported from multiple species and/or were far more common compared to others. In fact, Y132 was the most commonly reported substitution site by as many as 75 independent studies (Castanheira et al., 2020; Fan et al., 2023; Rao et al., 2024).

However, it is unclear if substitutions at these sites were truly imparting azole resistance or were preferentially identified and reported due to confirmation bias (Choi et al., 2018). At least, Y132 was known to interact with the ligand (Hargrove et al., 2017); and Y132F was shown to impart structural alteration by negating the hydrogen bond between the cofactor and the ligand, in turn leading to low binding efficiency between ERG11p and ligand (Paul et al., 2022).

Further, specific substitutions may impart resistance to some azoles but not to others. For instance, substitutions such as A114S, Y132F/H, K143Q/R, and Y257H were known to have much larger increase in MIC for fluconazole and voriconazole (≥4 fold) compared to itraconazole (≤2 fold) (Xiang et al., 2013). In addition, effects might vary widely between substitutions and azoles, and different substitutions in the same position might have different effects (Rao et al., 2024). Except for a few substitutions such as Y132F/H and K143R (Healey et al., 2018; Xiang et al., 2013), such information is unavailable for other substitutions and azole combinations. In fact, vast majority of the known substitutions were based on resistance to fluconazole, voriconazole, and “azole” with hardly any information on MIC values (Table S1). Thus, the literature information on resistant substitutions is by itself quite biased to short-tailed azoles and there is a need to explore them in the content of long-tailed azoles.

Based on a small number of substitutions, an early study observed that they all clustered in three regions that were then termed as hotspots (Marichal et al., 1999). Our comprehensive work found only about 48% of the sites within those previously recognized hotspot regions. While Y132F/H substitution that is at the azole-interacting site was known to have functionally deleterious effect (Hargrove et al., 2017; Paul et al., 2022; Rao et al., 2024), such information is scanty for other azole-interacting sites. We had hypothesized that the resistant substitutions should be disproportionately likely at the azole-binding sites (Rosam et al., 2021). However, that seemed to be false as just 33% of the known azole-interacting residues had known resistant substitutions, which is not very different from the expected. Further, very few instances of these substitutions/sites were reported. Mechanistic effect, if any, of substitutions occurring at non-azole-interacting sites remains to be elucidated (Rosam et al., 2021). It should also be noted that many substitutions are conservative and thus their effects, if any, are uncertain.

It is important to note that there were numerous reports of known “resistant” substitutions in azole sensitive/susceptible isolates (Asadzadeh et al., 2017; Liu et al., 2015; Rizzato et al., 2018; Silva et al., 2016; Table S3). We infer that such instances might be extremely common given the fact that sensitive isolates are seldom sequenced. This leads to an important and interesting question – do “resistant” substitutions really impart resistance? While a few might be as in Y132F/H (Hargrove et al., 2017; Paul et al., 2022; Rao et al., 2024), we argue that most substitutions are just sequence variations and might be unlikely to impart resistance at least on their own (however, they might impart resistance in combination with other substitutions). Azole resistance is complex and involves numerous mechanisms including mutations in other genes and the over expression of target genes and efflux pumps (Bhattacharya et al., 2020; Li et al., 2021; Rosam et al., 2021). This highlights the complexity of azole resistance and shows the need for further exploration to understand the functional significance of substitutions in the sterol 14α-demethylase enzyme.

To list some of the limitations of this work, we have collected the substitution information from the published literature, veracity of which might have indirectly affected our analyses and results. Being purely a computational study, we do not make any experimental validation as it is beyond the scope of this work.

In conclusion, we curated 2,222 instances of azole-resistant substitutions from *Candida* sterol 14α-demethylases. This will by itself serve as a useful reference catalogue for such studies in the future. We also presented relevant patterns such as site-specific abundance, and showed that substitutions did not occur disproportionately at azole-binding sites. Finally, we point to the potential bias that might exist in the data on azole-resistant substitutions. The results have relevance in the context of azole resistance and antifungal drug development.

## Funding and Acknowledgments

This work did not receive any specific funding.

## Statement of Ethics

The work is in compliance with ethical standards. No ethical clearance was necessary.

## Data Availability

The data used in this work were obtained from the literature. The relevant derived data are given in the supplemental tables.

## Author Contributions

RSPR and SDG planned the work. LP, SDG, and RSPR curated the data. RSPR analysed the data and wrote the paper. RPS, TCD, PNS, and VKS provided inputs for analysis/writing. All authors contributed intellectually, and edited/reviewed the manuscript. All authors have read and agreed to the published version of the manuscript.

## Supplemental Information

Supplemental information for this article is available online.

