## Supplemental for "Azole resistance: Patterns of amino acid substitutions in *Candida* sterol 14α-demethylase"

*Corresponding authors

^+^Present address

Running title: Substitution patterns in sterol 14α-demethylase

*Keywords: Amino acid substitution, Antifungal resistance, Azole binding, Candida infection, Computational sequence analysis*

**Table S1.** List of clinically/experimentally known amino acid substitutions imparting azole resistance in *Candida* sterol 14α-demethylase (Table_S1.xlsx).

**Table S2.** Azole-interacting residues (highlighted in dark green) in sterol 14α-demethylase based on PDB 3D structures.

|  | **4ZE3** | **5ESE** | **5ESM** | **7RY8** | **7RY9** | **7RYB** | **5TZ1** | **5UL0** | **7RYX** | **5FSA** | **5UL0** | **5ESG** | **5JLC** | **5V5Z** |  |
| --- | --- | --- | --- | --- | --- | --- | --- | --- | --- | --- | --- | --- | --- | --- | --- |
| **#** | **Flu** | **Flu** | **Flu** | **Vor** | **Vor** | **Vor** | **VT1** | **VT1** | **VT-1129** | **Pos** | **Pos** | **Itr** | **Itr** | **Itr** |  |
|  | **Sc** | **Sc** | **Sc** | **Sc** | **Sc** | **Sc** | **Ca** | **Sc** | **Sc** | **Ca** | **Sc** | **Sc** | **Cg** | **Ca** |  |
| 58 | V | V | V | V | V | V | F | V | V | F | V | V | V | F | . |
| 61 | A | A | A | A | A | A | A | A | A | A | A | A | A | A | * |
| 62 | V | V | V | V | V | V | A | V | V | A | V | V | I | A |  |
| 64 | Y | Y | Y | Y | Y | Y | Y | Y | Y | Y | Y | Y | Y | Y | * |
| 65 | G | G | G | G | G | G | G | G | G | G | G | e | G | G | * |
| 66 | M | M | M | M | M | M | Q | M | M | Q | M | M | T | Q |  |
| 87 | L | L | L | L | L | L | L | L | L | L | L | L | L | L | : |
| 88 | L | L | L | L | L | L | L | L | L | L | L | L | L | L | * |
| 118 | Y | Y | Y | Y | Y | Y | Y | Y | Y | Y | Y | Y | Y | Y | * |
| 121 | L | L | L | L | L | L | L | L | L | L | L | L | L | L | * |
| 122 | T | T | T | T | T | T | T | T | T | T | T | T | T | T | * |
| 126 | F | F | F | F | F | F | F | F | F | F | F | F | F | F | * |
| 131 | I | I | I | I | I | I | I | I | I | I | I | I | I | I | * |
| 132 | h | Y | Y | h | Y | h | Y | Y | Y | Y | Y | Y | Y | Y | * |
| 228 | F | F | F | F | F | F | F | F | F | F | F | F | F | F | * |
| 230 | P | P | P | P | P | P | P | P | P | P | P | P | P | P | * |
| 231 | I | I | I | I | I | I | I | I | I | I | I | I | I | I | : |
| 233 | F | F | F | F | F | F | F | F | F | F | F | F | F | F | * |
| 234 | V | V | V | V | V | V | V | V | V | V | V | V | V | V | * |
| 300 | L | L | L | L | L | L | L | L | L | L | L | L | L | L | * |
| 303 | G | G | G | G | G | G | G | G | G | G | G | G | G | G | * |
| 304 | V | V | V | V | V | V | I | V | V | I | V | V | V | I | : |
| 306 | M | M | M | M | M | M | M | M | M | M | M | M | M | M | * |
| 307 | G | G | G | G | G | G | G | G | G | G | G | G | G | G | * |
| 308 | G | G | G | G | G | G | G | G | G | G | G | G | G | G | * |
| 311 | T | T | T | T | T | T | T | T | T | T | T | T | T | T | * |
| 376 | L | L | L | L | L | L | L | L | L | L | L | L | L | L | * |
| 377 | H | H | H | H | H | H | H | H | H | H | H | H | H | H | * |
| 378 | S | S | S | S | S | S | S | S | S | S | S | S | S | S | * |
| 379 | L | L | L | L | L | L | I | L | L | I | L | L | L | I | : |
| 380 | F | F | F | F | F | F | F | F | F | F | F | F | F | F | * |
| 470 | C | C | C | C | C | C | C | C | C | C | C | C | C | C | * |
| 505 | F | F | F | F | F | F | Y | F | F | Y | F | F | F | Y | : |
| 506 | T | T | T | T | T | T | S | T | T | S | T | T | T | S |  |
| 507 | S | S | S | S | S | S | S | S | S | S | S | S | S | S | * |
| 508 | M | M | M | M | M | M | M | M | M | M | M | M | M | M | * |
| 509 | V | V | V | V | V | V | V | V | V | V | V | V | V | V | * |

Note: First row – PDB IDs; second row: ligand – short-tailed (fluconazole and voriconazole, light green shaded), medium-tailed (VT1 and VT-1129, light blue shaded), and long-tailed (posaconazole and itraconazole, light red shaded) azoles; third row: species – Ca (*C. albicans*), Cg (*C. glabrata*), and Sc (*S. cerevisiae*). First column is sequence index based on *C. albicans* and the last column is sequence conservation based on multiple sequence alignment of seven *Candida* sterol 14α-demethylase sequences. The protein-azole complexed 3D structures are not available for other combinations or *Candida* species.

**Table S3.** Example list of “resistant” substitutions (with more than 10 instances) in azole-sensitive/susceptible *Candida* isolates.

| **#** | **Substitution^1^** | **Species** | **# of instances** | **Reference** |
| --- | --- | --- | --- | --- |
| 1 | **D116E** | *C. albicans* | **21** | Ying et al., 2013 |
|  |  |  | **9** | Liu et al., 2015 |
|  |  |  | **7** | Xu et al., 2008 |
|  |  |  | **6** | Chau et al., 2004 |
|  |  |  | **5** | Xiang et al., 2013 |
|  |  |  | **2** | Manastır et al., 2011 |
|  |  |  | **1** | Sanglard and Bille, 2002 |
| 2 | **V437I** | *C. albicans* | **17** | Liu et al., 2015 |
|  |  |  | **9** | Xiang et al., 2013 |
|  |  |  | **4** | Ying et al., 2013 |
|  |  |  | **2** | Chau et al., 2004 |
|  |  |  | **1** | Manastır et al., 2011 |
| 3 | **E266D** | *C. albicans* | **14** | Ying et al., 2013 |
|  |  |  | **5** | Manastır et al., 2011 |
|  |  |  | **3** | Chau et al., 2004 |
|  |  |  | **3** | Xu et al., 2008 |
|  |  |  | **2** | Marichal et al., 1999 |
|  |  |  | **1** | Liu et al., 2015 |
|  |  |  | **1** | Xiang et al., 2013 |
| 4 | **R398I** | *Candida parapsilosis* | **8** | Arastehfar et al., 2020b |
|  |  |  | **6** | Kim et al., 2022 |
|  |  |  | **2** | Asadzadeh et al., 2017 |
|  |  |  | **2** | Castanheira et al., 2020 |
|  |  |  | **2** | Choi et al., 2018 |
|  |  |  | **1** | Berkow et al., 2015 |
| 5 | **K128T** | *C. albicans* | **6** | Ying et al., 2013 |
|  |  |  | **3** | Chau et al., 2004 |
|  |  |  | **3** | Xu et al., 2008 |
|  |  |  | **2** | Liu et al., 2015 |
|  |  |  | **1** | Manastır et al., 2011 |
|  |  |  | **1** | Sanglard and Bille, 2002 |
| 6 | **A114S** | *C. albicans* | **14** | Ying et al., 2013 |
| 7 | **Y257H** | *C. albicans* | **14** | Ying et al., 2013 |
| 8 | **Y132H** | *C. albicans* | **10** | Ying et al., 2013 |
|  |  |  | **2** | Liu et al., 2015 |
| 9 | **V488I** | *C. albicans* | **7** | Ying et al., 2013 |
|  |  |  | **3** | Chau et al., 2004 |
|  |  |  | **1** | Xu et al., 2008 |
|  | **…** | *…* | **…** | … |
|  | **30 substitutions (29 sites)^2^** | Five species | **258 instances** | 20 references |

^1^Site index is based on *C. albicans* sequence; ^2^See Table S1 for all such “resistant” substitutions in sensitive isolates.

```
albicans           ---------MAIVETVIDGINYFLSLSVTQQISILLGVPFVYNLVWQYLYSLRKDRAPLV	51
auris              ---------MALKDCIVDVVDRFSALPVPVKLAVLILVPIVYNLVWQFVYSLRKDRAPLV	51
glabrata           MSTENTSLVVELLEYVKLGLSYFQALPLAQRVSIMVALPFVYTITWQLLYSLRKDRPPLV	60
krusei             -MSVIKAIAADVQRYVLLAYSHFQTFSLLQQTLLVISIPFLYSALWQLLYSFRKDRVPMV	59
orthopsilosis      ---------MALVDLALQGYNYFMTLSTLQQFGLLVFAPFIYNIVWQLFYSLRKDRVPLV	51
parapsilosis       ---------MALVDLALHGYNYFMTLSTLQQFGLLVFAPFIYNIIWQLLYSLRKDRVPLV	51
tropicalis         ---------MAIVDTAIDGINYFLSLSLTQQITILVVFPFIYNIAWQLLYSLRKDRVPMV	51
                              :        . * ::    :  :::  *::*.  ** .**:**** *:*

albicans           FYWIPWFGSAASYGQQPYEFFESCRQKYGDVFSFMLLGKIMTVYLGPKGHEFVFNAKLSD	111
auris              FHWVPWVGSAVVYGMQPYQFFESCREKYGDVFAFVMLGKVMTVYLGPKGHEFVLNAKLAD	111
glabrata           FYWIPWVGSAIPYGTKPYEFFEDCQKKYGDIFSFMLLGRIMTVYLGPKGHEFIFNAKLAD	120
krusei             HYWIPWVGSAVVYGMQPYEFFENCRKQHGDVFSFLLLGKVMTVYLGPKGHEFVLNAKLSD	119
orthopsilosis      FYWIPWVGSAVSYGQDPYGFFEQCREKYGDLFAFVMLGRVMTVYLGPKGHEFVFNAKLSD	111
parapsilosis       FYWIPWVGSAVSYGQDPYGFFEQCREKYGDLFSFVMLGRVMTVYLGPKGHEFVFNAKLSD	111
tropicalis         FYWIPWFGSAASYGMQPYEFFEKCRLKYGDVFSFMLLGKVMTVYLGPKGHEFIYNAKLSD	111
                   .:*:**.***  ** .** ***.*: ::**:*:*::**::************: ****:*

albicans           VSAEDAYKHLTTPVFGKGVIYDCPNSRLMEQKKFAKFALTTDSFKRYVPKIREEILNYFV	171
auris              VSAEAAYSHLTTPVFGKGVIYDCPNSRLMEQKKFAKTALTKEAFQRYVPRIQEEVLDYFK	171
glabrata           VSAEAAYSHLTTPVFGKGVIYDCPNHRLMEQKKFVKGALTKEAFVRYVPLIAEEIYKYFR	180
krusei             VSAEDAYTHLTTPVFGKGVIYDCPNWKLMEQKKFAKVALTKESFIRYVPLIKDEMLKYFN	179
orthopsilosis      VSAEDAYQHLTTPVFGKGVIYDCPNARLMEQKKFAKTALTTDSFRRYVPLIRGEILDYFN	171
parapsilosis       VSAEDAYQHLTTPVFGKGVIYDCPNARLMEQKKFAKTALTTDSFRRYVPLIRGEILDYFT	171
tropicalis         VSAEEAYTHLTTPVFGKGVIYDCPNSRLMEQKKFAKFALTTDSFKTYVPKIREEVLNYFV	171
                   **** ** ***************** :*******.* ***.::*  *** *  *: .**

albicans           TDESFKLKEKTHGVANVMKTQPEITIFTASRSLFGDEMRRIFDRSFAQLYSDLDKGFTPI	231
auris              ACSQFKMNERNNGVANVMKTQPEMTILTASKSLMGDDMRARFDASFAKLYSDLDKGFTPI	231
glabrata           NSKNFKINENNSGIVDVMVSQPEMTIFTASRSLLGKEMRDKLDTDFAYLYSDLDKGFTPI	240
krusei             ANF-----RGDSGKTDVLKSQSEMTLFTASRSLFGDALRNRLDASYAEMYSDLDKGFTPL	234
orthopsilosis      KSKVFNMKTKKSGVVDVLQSQPEITIFTASRSLLGEAMRQRFDASFAQLYADLDKGFTPI	231
parapsilosis       KSKVFNMKKQKSGVVDVLQSQPEITIFTASRSLLGEAMRKRFDASFAQLYADLDKGFTPI	231
tropicalis         NDVSFKTKERDHGVASVMKTQPEITIFTASRCLFGDEMRKSFDRSFAQLYADLDKGFTPI	231
                               * ..*: :* *:*::***:.*:*. :*  :* .:* :*:********:

albicans           NFVFPNLPLPHYWRRDAAQKKISATYMKEIKSRRERGDIDPNRDLIDSLLIHSTYKDGVK	291
auris              NFVFPHLPLPAYWKRDAAQQKISATYMSLINERRKTGDIVPDRDLIDSLMTNSTYKDGVK	291
glabrata           NFVFPNLPLEHYRKRDHAQQAISGTYMSLIKERREKNDIQ-NRDLIDELMKNSTYKDGTK	299
krusei             NFVFSYLPLPNYWKRDAAHKNISNTYLDLINTKRAGGEIK-NEDLVDALLKNSVYKDGTR	293
orthopsilosis      NFVFPHLPLPHYWKRDAAQQKISETYMKEIARRRESGDIDENRDLIDSLLVNSTYKDGVK	291
parapsilosis       NFVFPHLPLPHYWKRDAAQQKISETYMTEIARRRETGDIDENRDLIDSLLVNSTYKDGVK	291
tropicalis         NFVFPNLPLPHYWRRDAAQRKISAHYMKEIKRRRESGDIDPKRDLIDSLLVNSTYKDGVK	291
                   ****  ***  * :** *:: **  *:  *  :*  .:*  ..**:* *: :*.****.:

albicans           MTDQEIANLLIGILMGGQHTSASTSAWFLLHLGEKPHLQDVIYQEVVELLKEKGGDLNDL	351
auris              MTDQEVANLLIGVLMGGQHTSASTSAWFLLHLAEQPKLQEELYNEVLSVLAEKGGSLKDL	351
glabrata           MTDQEIANLLIGVLMGGQHTSAATSAWCLLHLAERPDVQEELYQEQMRVLNN---DTKEL	356
krusei             MTDEELAHLMIGVLMGGQHTSSATSAWFLLHLGEKPQLQEEIYREIQSVLGEN--FEREL	351
orthopsilosis      MTDQEIANLLIGVLMGGQHTSATTSAWFLLHLAEKPQLQDELYQEVLNALSGKGGNLDDL	351
parapsilosis       MTDQEIANLLIGVLMGGQHTSATTSAWFLLHLAEKPQLQDELYQEVLNALSGKGGNLDDL	351
tropicalis         MTDQEIANLLIGVLMGGQHTSASTSAWFLLHLAEQPQLQDDLYEELTNLLKEKGGDLNDL	351
                   ***:*:*:*:**:********::**** ****.*:*.:*: :*.*    *        :*

albicans           TYEDLQKLPSVNNTIKETLRMHMPLHSIFRKVTNPLRIPETNYIVPKGHYVLVSPGYAHT	411
auris              AYDDLQKMPLINQTIKETLRLHMPLHSIFRKVMNPLVVPNTKYVVPKGHYVMVSPGYAQT	411
glabrata           TYDDLQNMPLLNQMIKETLRLHHPLHSLFRKVMRDVAIPNTSYVVPRDYHVLVSPGYTHL	416
krusei             TYDDLQKLDLVNATIKETLRLHMPLHSIFRKVTRDLPVPNTSYIVPKGHYVLISPGYTML	411
orthopsilosis      SYEDLQQMPLVNNTIKETLRLHMPLHSIFRKVVSPLVVPNTKYIVPKGHHVLVSPGYAHT	411
parapsilosis       SYEDLQQMPLVNNTIKETLRLHMPLHSIFRKVVSPLVVPNTKYIVPRGHHVLVSPGYAHT	411
tropicalis         TYEDLQKLPLVNNTIKETLRMHMPLHSIFRKVMNPLRVPNTKYVIPKGHYVLVSAGYAHT	411
                   :*:***::  :*  ******:* ****:****   : :*:*.*::*:.::*::* **:

albicans           SERYFDNPEDFDPTRWDTAAAKANSVS-----FNSSDEVDYGFGKVSKGVSSPYLPFGGG	466
auris              NEKWFPRANEFDPHRWDEETSS----------NIDTDAVDYGFGKVTKGVSSPYLPFGGG	461
glabrata           QEEFFPKPNEFNIHRWDGDAASSS--------AAGGDEVDYGFGAISKGVSSPYLPFGGG	468
krusei             SERYFPNASEFQPHRWDEIKSIDGGISLPAEGENAKETVDYGFGKISKGVASPYLPFGGG	471
orthopsilosis      NERFYKDASAFNPHRWDESAS-----------TNDAGEVDYGFGKVSKGVSSSYLPFGGG	460
parapsilosis       NERFYKDASDFNPHRWDESAS-----------TNDAGEVDYGFGKVSKGVSSSYLPFGGG	460
tropicalis         SDRWFEHPEHFNPRRWESDDTKASAVS-----FNSEDTVDYGFGKISKGVSSPYLPFGGG	466
                   .:.::   . *:  **:   :                 ****** ::***:* *******

albicans           RHRCIGEQFAYVQLGTILTTFVYNLRWTIDG-Y-KVPDPDYSSMVVLPTEPAEIIWEKRE	524
auris              RHRCIGEQFAYVQLGTILATYVYNIKWRFKKDG-SLPPVDYQSMVTLPMEPAEIEWEKRE	520
glabrata           RHRCIGELFAYCQLGVLMSIFIRTMKWRYPTEGETVPPSDFTSMVTLPTAPAKIYWEKRH	528
krusei             RHRCIGEPFAYTQLGTLLVHYIQNFKWTA-----KVPPIDYTSMVTLPTQPAEIKWEGRQ	526
orthopsilosis      RHRCIGEQFAYVQLGTILTTFVYNVKWKLAN-G-KVPDVDYTSMVTLPQDPAEIVWEKRD	518
parapsilosis       RHRCIGEQFAYVQLGTILTTFVYNLKWKLAN-G-KVPDVDYTSMVTLPQHPAEIVWEKRD	518
tropicalis         RHRCIGEQFAYVQLGTILTTYIYNFKWRLNG-D-KVPDVDYQSMVTLPLEPAEIVWEKRD	524
                   ******* *** ***.::  :: ..:*       .:*  *: ***.**  **:* ** *.

albicans           TCMF-	528
auris              TCVY-	524
glabrata           PEQKY	533
krusei             KN---	528
orthopsilosis      TCVL-	522
parapsilosis       TCVI-	522
tropicalis         TCMV-	528
```

**Fig. S1.** Multiple sequence alignment of sterol 14α-demethylase proteins from seven *Candida* species (*C. albicans* [UniProt ID: P10613], *C. auris* [A0A2H1A309], *C. glabrata* [P50859], *C. krusei* – currently renamed as *Pichia kudriavzevii* [Q09GQ6], *C. orthopsilosis* [H8X874], *C. parapsilosis* [C7EXA5], and *C. tropicalis* [P14263]). The multiple sequence alignment was done using the Clustal Omega web server (https://www.ebi.ac.uk/Tools/msa/clustalo/, last accessed on Apr 24, 2024). Kwon substitution sites are blue shaded.
